# High overexploitation risk due to management shortfall in highly traded requiem sharks

**DOI:** 10.1101/2022.06.09.495558

**Authors:** C. Samantha Sherman, Eric D. Digel, Patrick Zubick, Jonathan Eged, Alifa B. Haque, Jay H. Matsushiba, Colin A. Simpfendorfer, Glenn Sant, Nicholas K. Dulvy

**Affiliations:** Earth to Oceans Research Group, Department of Biological Sciences, Simon Fraser University, Burnaby, British Columbia, Canada; TRAFFIC International, Cambridge, United Kingdom; Nature-based Solutions Initiative, Department of Zoology, University of Oxford, UK; Department of Zoology, University of Dhaka, Bangladesh; Institute of Marine and Antarctic Studies, University of Tasmania, Hobart, Tasmania, Australia; Australian National Centre for Ocean Resources and Security, University of Wollongong, NSW, Australia

**Keywords:** Carcharhinidae, CITES, fisheries management, international policy, international trade, Shark conservation, shark fin trade, shark meat

## Abstract

Most of the international trade in fins (and likely meat too) is derived from requiem sharks (family Carcharhinidae), yet trade in only two of the 56 species is currently regulated. Here, we quantify catch, trade, and the shortfall in national and regional fisheries management (M-Risk) for all 56 requiem shark species based on 831 assessments across 30 countries and four Regional Fisheries Management Organisations (RFMOs). Requiem sharks comprise over half (60%) of the annual reported global chondrichthyan catch with most species (86%) identified in the international fin trade. Requiem sharks are inadequately managed by fisheries, with an average M-Risk of half (50%) of an ideal score, consequently 70% of species are threatened globally. The high catch and trade volume and shortfall in management of these iconic species requires a global integrated improvement in fisheries management, supported by regulating international trade to sustainable levels.

## 1 INTRODUCTION

While there are promising signs that conservation is working in the oceans, overfishing remains the principal cause of biodiversity loss (Duarte et al. 2020). Fisheries targeting bony fishes are increasingly being brought into sustainability. For example, sustainable fisheries are found in regions and nations with high capacity for management like Regional Fisheries Management Organisations (RFMOs) and developed countries in North America, Europe as well as South Africa, Australia, and New Zealand (Simpfendorfer and Dulvy 2017, Hilborn et al. 2020). Yet, other species taken in these fisheries, notably sharks and rays, continue to decline in part because of their more sensitive life histories (Myers and Worm 2005), but also because of a lack of management (Juan-Jordá et al. 2017, Fordham et al. 2022). The catch of shark, ray, and chimaera (Class Chondrichthyes, hereafter ‘sharks’) is poorly monitored globally, with only one-third of reported catch identified beyond Family level (FAO 2022). Yet locally, sharks provide significant income for small-scale fishers and individual specimens are retained because of the extremely high value of their body parts, such as fins, jaws, skins, and liver oil in the international trade markets (McClenachan et al. 2016, Booth et al. 2021).

The latest comprehensive global Red List assessment of sharks reveals that one-third (37.5%) of all species are threatened with an elevated risk of extinction (Dulvy et al. 2021). Notably, many groups, such as tropical sharks and rays, are at very high risk of extinction (Dulvy et al. 2021). A key way to halt and reverse the declines of sharks is through quantifying and improving national fisheries management. Here, we focus on understanding national fisheries management of sharks that are highly threatened, in part due to international trade demand for meat and fins (Dent and Clarke 2015, Niedermüller et al. 2021).

The requiem sharks (family Carcharhinidae) are a recent radiation found throughout tropical and subtropical waters, which have historically dominated the shark catches of coastal fisheries (Lam and Sadovy de Mitcheson 2011). Indeed, this is one of seven priority shark taxa based on the high volume of fisheries catches taken in poorly documented and poorly regulated tropical multispecies fisheries (Dulvy et al. 2017). Further, many species of the family are highly prevalent in the international fin trade (Fields et al. 2017, Cardeñosa et al. 2020).

Here, we ask whether fisheries management is adequate to halt declines and improve the status of requiem sharks. We estimate the risk due to overexploitation based on the current state of global fisheries management for all 56 species from the family Carcharhinidae using a novel rapid assessment of management risk (M-Risk)(Sherman et al. 2022). Specifically, we ask three questions: (1) how prevalent are requiem sharks in global catch and international trade?, (2) are requiem sharks adequately managed across their geographic range? and which (if any) species are adequately managed?, and (3) which forms of management are most prevalent and which are lacking?

## 2 METHODS

### 2.1 Selection of management units

Country management units were selected based on their proportional contribution to global shark catch, (2010–2019, inclusive) (FAO 2022). We included the 21 countries that cumulatively report >80% of global Chondrichthyan catch in addition to nine countries with high catch in each FAO fishing region to ensure global representation. In addition to countries, all four tuna Regional Fisheries Management Organisations (RFMOs) were included as management units due to the high volume of shark catch in Areas Beyond National Jurisdictions (total management units = 34) (Tolotti et al. 2015, Murua et al. 2021). Our references to ‘management units’ refer to the jurisdictions and not specific stocks.

## 2.2 M-Risk: Management risk assessment

Species assessments were completed by scoring management risk against 21 measurable attributes, as fully explained in Sherman *et al*. (2022) and described in **Supplemental Material 1**. The 21 Attributes were split into five classes, three scored for all management units (Management System [*n*=5 Attributes], Fishing Practises & Catch [*n*=5], Compliance, Monitoring & Enforcement [*n*=5]), and two that were specific to either a country (*n*=4) or RFMO (*n*=2).

## 2.3 Scoring of Attributes

All attributes were scored in an ordinal manner with a narrow range of scores to ensure consistency (i.e., 0–3, 0–4, or 0–5) such that higher scores indicated management with a higher likelihood of sustainable outcomes (for full details see **Supplemental Material 1**). Assessments were completed through exhaustive internet searches and national fisheries department websites. Where English was not the official language, searches were completed in the official language and documents were translated using Google translate. To accommodate the potential for ambiguous translations, points were generously awarded. However, if no information for an attribute was found, a precautionary score of zero was given. The final ‘Management Score’ was expressed as a percentage of the total possible points. This score indicates progress towards the ideal management (of 100%), therefore, a score greater than 75% could be considered a well-managed fishery for sharks.

## 2.4 Additional Catch, Trade, and Species Data

Identification of Carcharhinidae species in international trade was obtained through a thorough literature search (**Supplementary Material 2**). To determine threat level of all Carcharhinidae, the IUCN Red List Status of all species and species distribution maps were sourced after the 2021-03 update of the IUCN Red List website (www.iucnredlist.org)(Dulvy et al. 2021). Shark catch from all countries and RFMOs was sourced from FishStatJ v4.02.07 (FAO 2022), and the Sea Around Us (www.seaaroundus.org)(Pauly et al. 2021). A generalised linear model (GLM) comparing volume of shark catch and mean management score was completed (**Supplementary Material 3**).

## 3 RESULTS

### 3.1 What is the scale of catch and trade of requiem sharks?

Based on the FAO global catch data from 2010–2019, inclusive, 65.5% of Chondrichthyan catch was reported only as Class (Chondrichthyes), subclass (Elasmobranchii), or superorder (Selachimorpha). Carcharhinidae species accounted for 59.6% of shark catch reported below Order, and 44.2% of reported coastal shark catch (excluding Blue Shark, *Prionace glauca*; 438,799 and 325,418 metric tonnes per year, respectively)(FAO 2022). Over three-quarters of Carcharhinidae species (minimum 80.4%, n = 45 species, to 85.7%, n = 48) have been documented in the international fin trade either through market sampling or shipment seizures (**Supplementary Material 2**). Based on the under-reporting of shark catch to the FAO by up to four times (Clarke et al. 2006), the true volume of Carcharhinidae catch is likely closer to 1,755,200 mt per year and possibly higher if the broad catch reporting includes Carcharhinids.

### 3.2 Are requiem sharks adequately managed across their geographic range?

Requiem sharks are inadequately managed worldwide, both by nations and RFMOs, and are globally distributed with hotspots in northern Australia and Southeast Asia (**Figure 1a**). Across 831 M-Risk assessments from 30 countries spanning all six inhabited continents and four RFMO fishing grounds, the average management risk was high, with only half of the ideal management in place for requiem sharks (50.0% ± 0.6, n = 831). Regionally, the management shortfall for requiem sharks was greatest in Africa (mean management score 36.3% ± 1.5, n = 88) and Asia (38.6% ± 0.4, n = 332; **Figures 1b and S1**). Both regions were much lower than the remaining regions and RFMO average: Oceania (67.1% ± 1.2, n = 122), Europe (62.9% ± 1.4, n = 21), North America (60.9% ± 1.1, n = 130), RFMOs (60.5% ± 0.7, n = 63), and South America (56.7% ± 0.7, n = 75) (**Figures 1b,c and S1**).

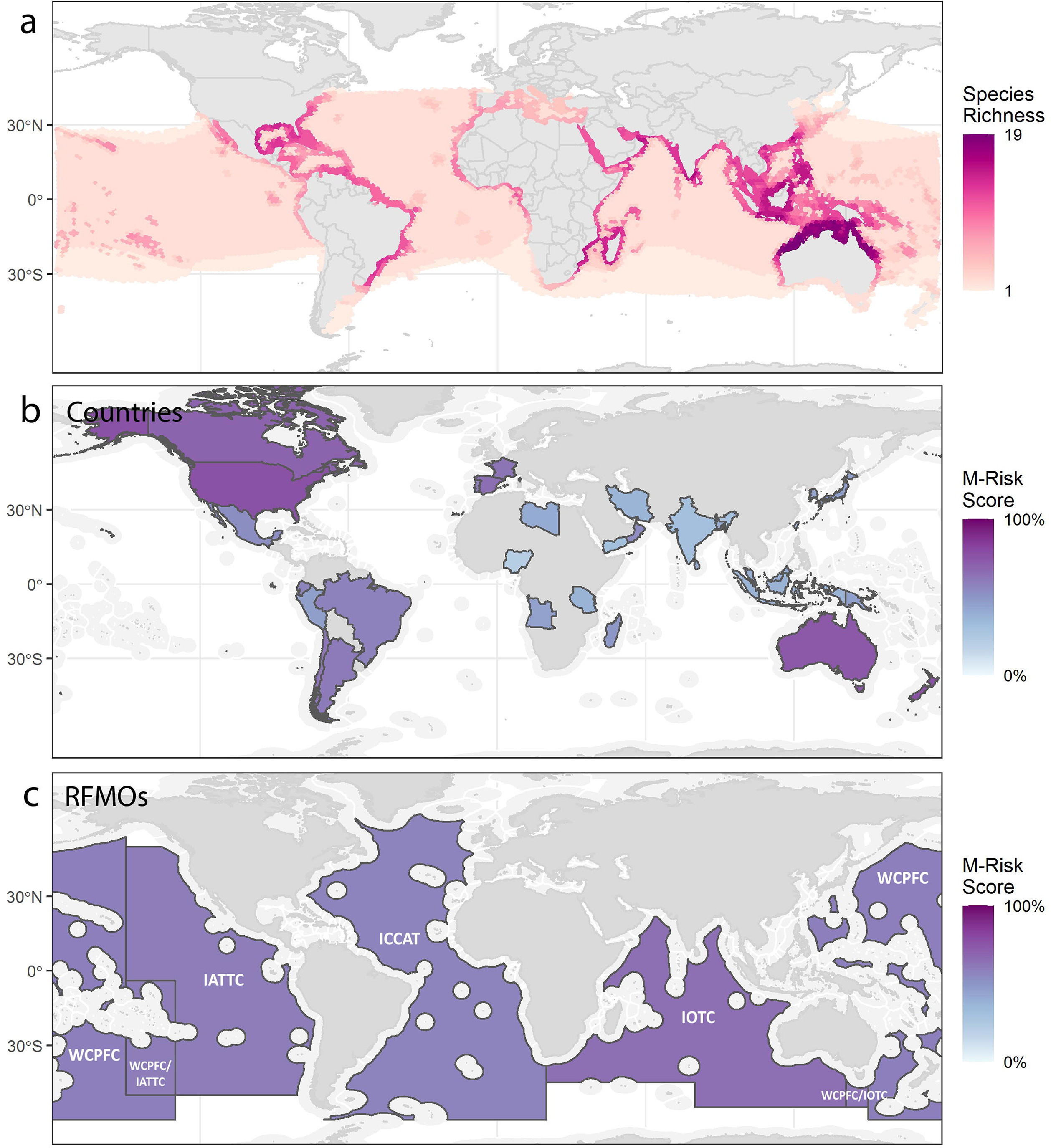

Some of the biggest fishing nations of the world have the lowest fisheries management scores including Nigeria (21.9% ± 0.0, n = 12), India (30.4% ± 0.4, n = 25), Yemen (31.5% ± 0.3, n = 20), and Iran (36.5% ± 0.6, n = 17) (**Figures 1b and 2**). These countries are all among the top 20 shark catching countries globally (Okes and Sant 2019). Much of their catch is exported through non-standardized commodity codes, masking catch details (Dent and Clarke 2015). The countries with the highest scores, and assessments on more than three species, were the USA (76.8% ± 1.2, n = 18), Australia (74.1% ± 0.5, n = 29), Spain (65.7% ± 2.2, n = 7), and Argentina (61.0% ± 1.0, n = 5), respectively (**Figures 1b and 2**). Interestingly, these countries represent four different continents, indicating there are ‘pockets’ with better management scores (above 60%) for requiem sharks. Three of these countries (all except Australia) also rank among the top 20 shark catching countries globally (Okes and Sant 2019), indicating catch quantity is not a direct result of management. Indeed, there was no relationships between volume of shark catch in each management unit and the respective management scores (GLM, t = 1.46, df = 32, p = 0.153; **Figure 2b**).

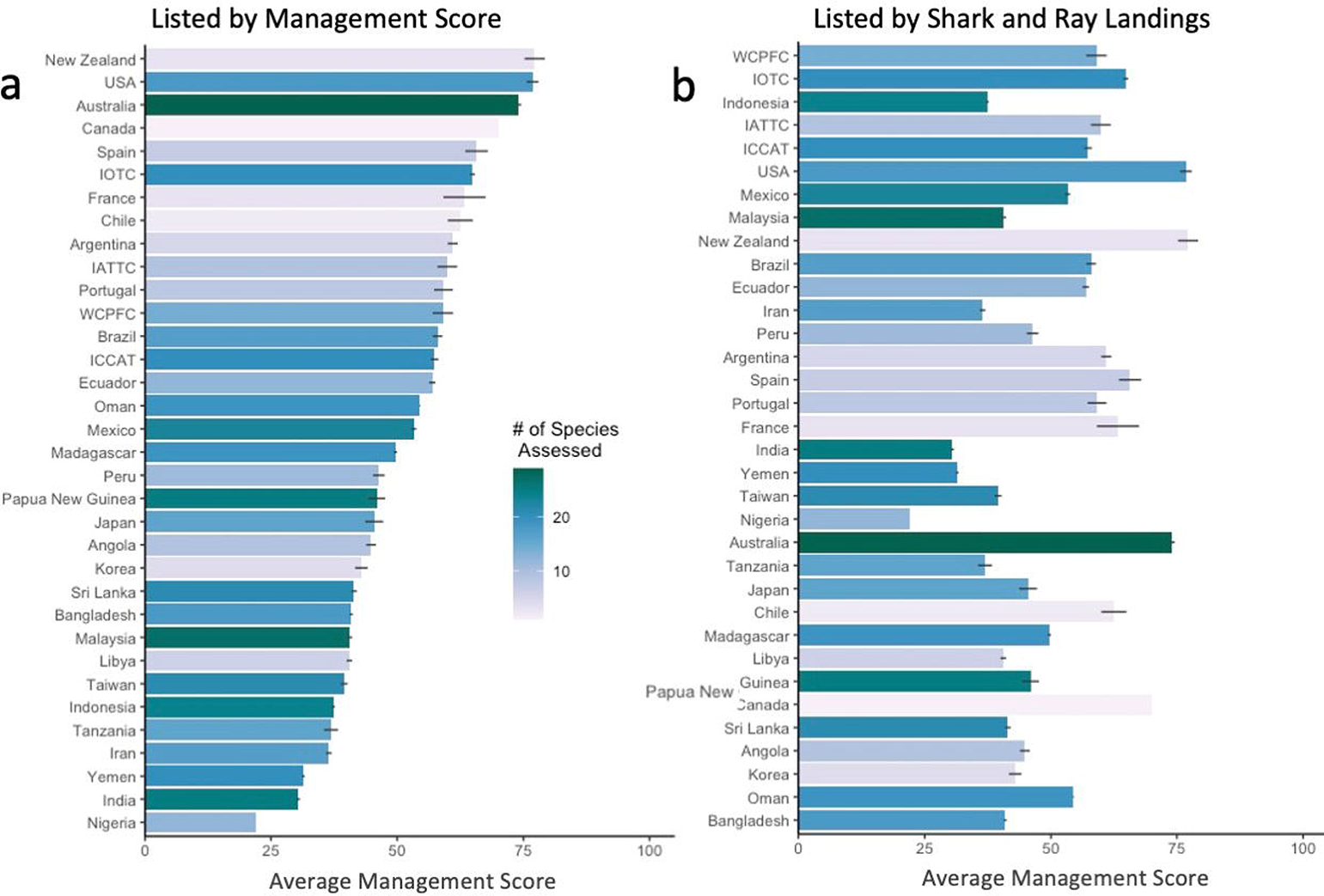

### 3.3 Which species are adequately managed?

Strictly speaking, any species with an M-Risk score of less than 100% is undermanaged. All species are undermanaged, by that criterion, across a total of 831 assessments. The species with the highest mean management score, assessed in three or more countries, are Blacknose Shark (*C. acronotus*: 62.4% ± 7.0, n = 4), Atlantic Sharpnose Shark (*Rhizoprionodon terraenovae*: 61.1% ± 7.4, n = 3) and Caribbean Reef Shark (*C. perezi*: 60.3% ± 6.6, n = 3) (**Figure 3**). These three species have relatively small distributions and occur in the Western Central and Southwest Atlantic Ocean. Status is a function not only of management but also total catch volume, therefore in regions with higher fishing effort and catch like the Western Atlantic (Bell et al. 2017), even higher management scores are not currently effective at ensuring sustainable catch levels for the species with slower life histories (Worm et al. 2013). Hence, the Blacknose Shark and Caribbean Reef Shark are both listed as Endangered (EN) as per the IUCN Red List and the Atlantic Sharpnose Shark is Least Concern (LC), in part because of its faster life history.

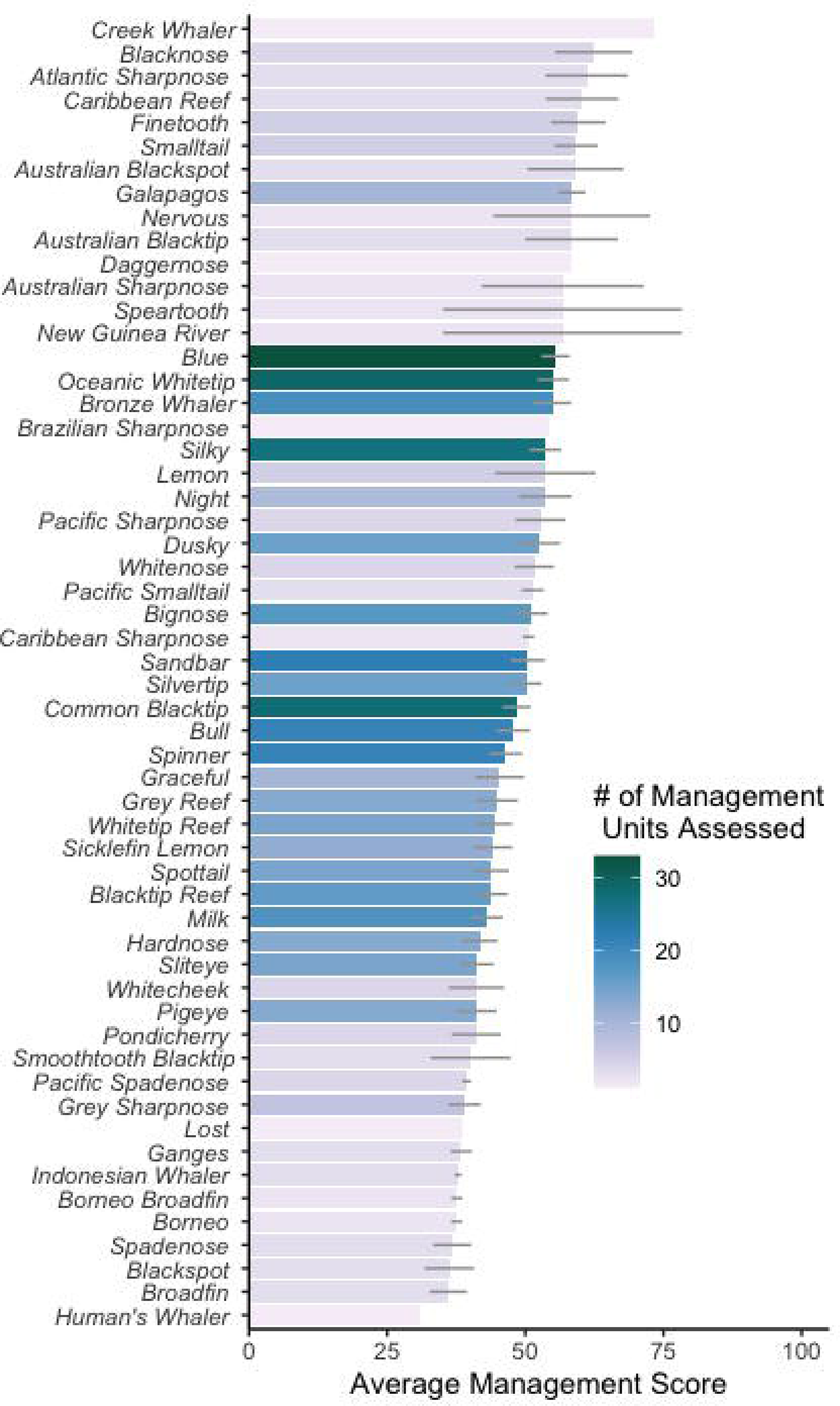

The lowest scoring species (i.e., the *least* managed), assessed in at least three countries, were the Broadfin Shark (*Lamiopsis temminckii*: 36.1% ± 3.4, n = 3), followed by the Blackspot Shark (*C. sealei*: 36.3% ± 4.5, n = 3), and Spadenose Shark (*Scoliodon laticaudus*: 36.8% ± 3.5, n = 3) (**Figure 3**). The first two are listed in threatened categories (EN and Vulnerable-VU, respectively), with the third being Near Threatened-NT, and all occur in the heavily fished waters of the Indian Oceans (Blaber et al. 2009, Pauly et al. 2021).

The variation between the single highest and lowest management score across all management units were 21.9% for all 13 species in Nigeria and 85.0% for Copper Shark (*C. brachyurus*) in the USA, indicating a wide variation in management for species (over 60%). Species with intermediate management scores tended to be assessed in a greater number of management units. For example, the Sandbar Shark (*C. plumbeus*: 50.5% ± 3.1), Silky Shark (*C. falciformis*: 53.6% ± 2.9), and Blacktip Shark (*C. limbatus*: 48.4% ± 2.6) were assessed in 22, 27, and 28 different management units, respectively, and are all listed as threatened (EN, VU, VU, respectively) (**Figure 3**). The three highest and lowest scoring species were only assessed in three or four management units, indicating potential regional areas of management concern (i.e., Northern Indian Ocean) or hope (i.e., Western Atlantic).

### 3.4 Which aspects of management are most prevalent and lacking?

Across all assessed management units, the attributes with the highest average scores were lower resolution structural attributes, related to general fisheries management, such as: if a regulatory body was present (95.7% ± 2.5), engagement with CITES (93.3% ± 2.8) or Ecosystem Based Fisheries Management (EBFM) for RFMOs (91.7% ± 2.9), followed by IUU management (84.8% ± 4.7) (**Figure 4a**). Attributes with lower average scores were the harder to implement, species-specific fisheries operational attributes including understanding of the species’ stock status (18.3% ± 4.9), species-specific compliance measures to reduce fishing mortality (21.0% ± 3.8), and taxonomic resolution of landing limits in place or if any limits existed (21.2% ± 4.5) (**Figure 4a**). These attribute scores indicate that while countries and RFMOs have the foundations in-place to incorporate management, this is not yet being directed towards significant catches of threatened species, like the requiem sharks. The pattern of RFMO attribute scoring of species was similar to management units, with greater scores for structural attributes and lower scores for fisheries operational attributes (**Figure 4b**). For some species, like the CITES-listed Oceanic Whitetip and Silky Sharks (*C. longimanus* and *C. falciformis*), landing limits existed more often, and at higher resolution than the average requiem shark (65.6% ± 8.5 and 49.6% ± 9.5, respectively, vs. 20.0% ± 2.9 average of all requiem sharks).

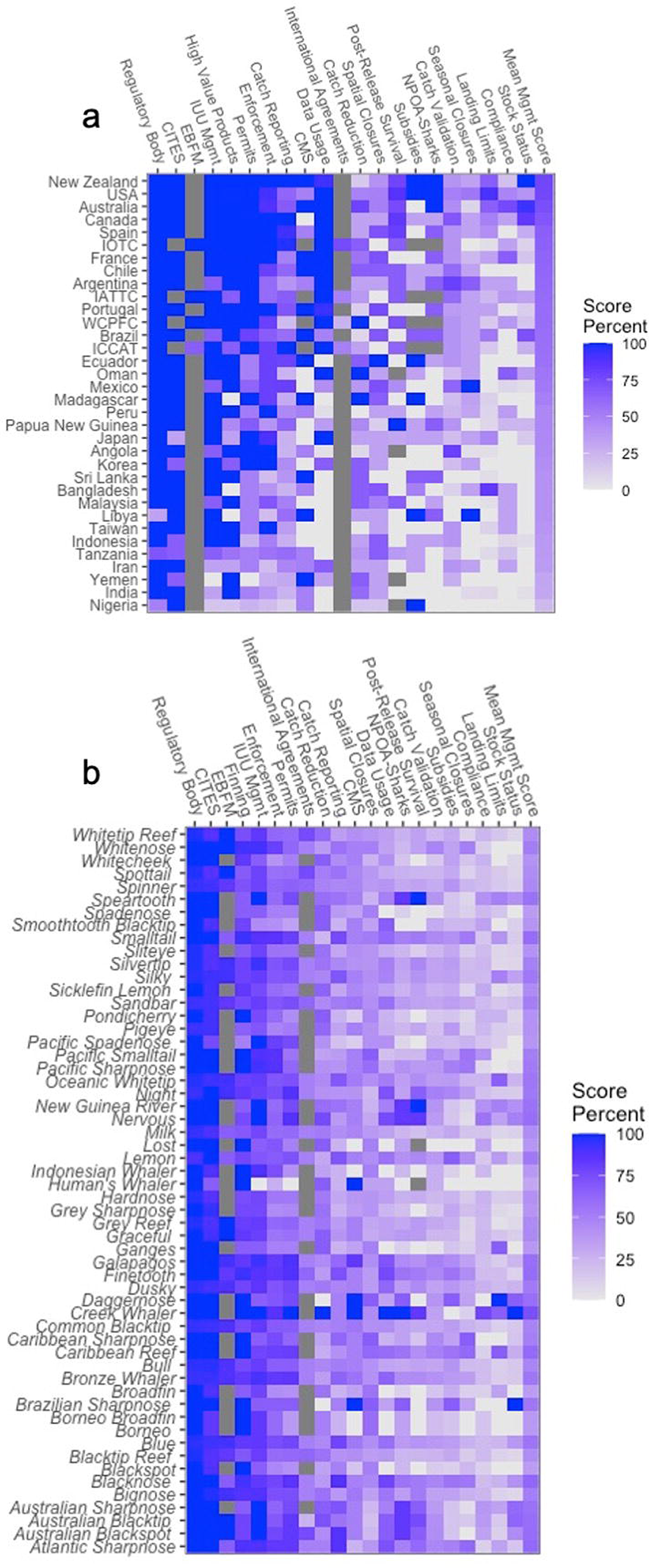

## 4 DISCUSSION

We have three key findings. First, the global catch of requiem sharks is almost two-thirds of the global reported catch of all sharks reported to a higher resolution than Order with a volume likely over 1.75 million metric tonnes per year. Requiem sharks are also highly represented in international trade, with up to 86% of species in the Carcharhinidae family identified. Second, the management risk scores for 56 species of requiem sharks from 30 nations and four RFMOs demonstrate they are undermanaged with only half of the ideal management in place across all species. M-Risk is lower in Africa and Asia and higher in Oceania, but there are better managed countries found on each continent, representing pockets of hope. Third, management deficits typically exist in species-specific regulations, while general fisheries structural attributes often scored higher. Taken together, these three catch, trade, and management shortfall issues have likely resulted in high levels of overfishing of the Carcharhinidae family inferred from an elevated risk of extinction. We next consider the following issues (1) how to improve domestic management, (2) improving global fisheries management, (3) the emerging opportunity that trade regulation presents, and (4) the benefits of trade regulation of requiem sharks.

Globally, requiem sharks have minimal species-specific fisheries regulations, including a lack of landing limits, and coarse catch reporting generalised to the genus, class, or often not reported at all. The recognition that many shark species, including requiem sharks, are threatened by overfishing, has resulted in increasing national and international efforts to improve their management. However, there is little evidence that these efforts have been able to address the issue in a significant way (Davidson and Dulvy 2017). In addition to a lack of domestic measures, documentation meant to ensure internationally traded species are caught in a sustainable manner (i.e., catch documentation schemes) are not used for requiem sharks (Mundy-Taylor et al. 2014). Domestic and international regulations have increased in the past decade to eliminate the removal of fins and discarding of carcases (shark finning) but unsurprisingly have had little effect on reducing the volumes of fins traded, because they do not limit fishing mortality (Lawson and Fordham 2018, Ferretti et al. 2020). In addition to the fin trade, there has been a recent increase in the international shark meat trade (Dent and Clarke 2015, Liu et al. 2021). This may be adding to demand and, therefore, incentive for fishers to land sharks, thus additional constraints are urgently needed to ensure catches are sustainable. Management units with higher shark catch volumes are not necessarily better managed, but can be used to provide priority areas for management improvement (e.g., Indonesia, which has high shark catch but a lower management score).

Globally widespread species of sharks are subject to fragmented management, requiring a coherent response (Dulvy et al. 2017). The wide-ranging requiem sharks suffer from a patchwork of management and might have lower extinction risk only if a significant part of their range lies within the waters of better managed countries like the USA and Australia, e.g., Blacktip Reef Shark (*C. melanopterus*), which is well-managed in Australia, but less so elsewhere in its range (**Figures 1 and 2**). The broad distributions of many species, and large volume of requiem sharks in global trade means the need for well-managed fisheries is pertinent to all range states to combat the problem of trade-driven ‘roving bandit dynamics’ (Berkes et al. 2006). Traded shark products often travel through several different countries being aggregated, processed, and then reimported for sale to the general public resulting in poor traceability and species identification and hence ability understand the role of trade in driving local mortality (Haque and Spaet 2021, Niedermüller et al. 2021). The onus should be on both importers and exporters to ensure traded products are fished sustainably and legally acquired. Improving management and introducing national and/or fishery landing limits for requiem sharks would be an essential part of ensuring sustainable responsible fisheries. However, this must be completed coherently in all countries to ensure future sustainability of requiem shark catches, and this could be achieved through international trade regulation. There is an immediate need for global cooperation in the monitoring, sustainable catch, and trade of these species, which can be achieved through international agreements such as CITES, CMS, and adherence to the FAO Guidelines for Responsible Fish Trade.

Listing species on Appendix II of CITES ensures that specimens cannot be traded unless they are legally acquired from a sustainable fishery (i.e., have a positive Non-Detriment Finding, NDF), and both exporting and importing countries have permits (Fernando et al. 2022). Hence, such a listing would incentivise assessment and management of national fisheries. The development of tools to assess whether a fishery is detrimental or otherwise has driven investment in monitoring and management of CITES Appendix II listed sharks (Martin 2007). There are promising signs of progress, even though measuring effectiveness is challenging, and the listings have been comparatively recent relative to the long generation lengths and recovery times of sharks (Friedman et al. 2018). Appendix II CITES listings have led to detectable progress in governance, particularly in major shark fishing nations like Indonesia and Malaysia (Friedman et al. 2018). In Indonesia, implementation of listings resulted in reduced catch and trade of manta rays (*Mobula alfredi* and *M. birostris*)(Booth et al. 2020). Here, we identify higher scores in species-specific attributes, like landing limits, for the CITES-listed Oceanic Whitetip and Silky Sharks (*C. longimanus* and *C. falciformis*). Requiem shark trade from Indonesia (the top shark catching country in the world) is reported to Family-level and contributes 51% to their shark production (Prasetyo et al. 2021). This broad level of reporting may be masking serial depletions. Listing all species together would minimise enforcement challenges and avoid the complex challenges of separating individual requiem shark species by customs officials in shipments (Partin et al. 2022). However, both obligations of a CITES listing: (1) to have species-specific positive NDFs, and (2) to report species-specific trade data, would greatly improve our ability to discern species-specific fishing mortality using trade volume data (Janssen and Leupen 2019). Evidence of improved species status after CITES listings is minimal, but has been observed for mammals and it is incontrovertible that many extinctions have been avoided because of CITES (Mialon et al. 2022). However, in some cases, anticipation of CITES listing and ratification may spur traders to increase trade volume prior to the need for an NDF (Mialon et al. 2022). Despite this initial increase, regulations moving forward would allow for species control and recovery, therefore, long-term sustainability is only possible through immediate action.

### Policy relevance

Requiem sharks are heavily fished and traded, with at least 44, and up to 48 species of the family Carcharhinidae documented in international markets and fin seizures (**Supp. Data S2**). Most requiem sharks are threatened and indeed the Carcharhinidae family is in the top ten most threatened families of sharks and rays, with over two-thirds (69.6%, 39 of 56) of species threatened (Dulvy et al. 2021). Here, we show that this high level of threat, from overfishing and unregulated international trade, results from a shortfall in national and international management. The two CITES listed requiem sharks have higher scores in species-specific attributes, showing the effectiveness of CITES listing on management. We, therefore, conclude that listing requiem sharks on CITES Appendix II provides a coherent mechanism for tackling the worldwide management deficits for requiem sharks.

## ACKNOWLEDGEMENTS

The authors thank Brittany Finucci for her comments on study design. We also thank the following people for help finding or understanding legislation in various countries: Maite Pons (Argentina), David Kulka (Canada), Sushmita Mukherji and Zoya Tyabji (India), Aiko Matsushiba (Japan), Sara A.A. Al Mabruk (Libya), Gonzo Araujo and Cat McCann (Malaysia), Brittany Finucci (New Zealand), Leontine Baje and Michael Grant (Papua New Guinea), Andrew Bickell (Sri Lanka), Crystal McRae (Taiwan), and Tobey Curtis and John Carlson (USA). This work was supported by a grant to GS from the Shark Conservation Fund, a philanthropic collaborative that pools expertise and resources to meet the threats facing the world’s sharks and rays. The Shark Conservation Fund is a project of Rockefeller Philanthropy Advisors. NKD was supported by the Discovery and Accelerator grants from Natural Science and Engineering Research Council and the Canada Research Chair program. The authors have no conflicts of interest to declare.

## Notes

### Competing Interest Statement

The authors have declared no competing interest.

### Summary of Updates

Minor edits upon resubmission to journal

